# High-density lipoprotein mimetic peptide 4F ameliorates APOE4-associated lipid dysfunction in primary and iPSC-derived astrocytes and cerebral organoids

**DOI:** 10.64898/2025.12.16.694774

**Authors:** Kristina Fredriksen, Siddhi S. Joshi, Allison Chang, Ling Li

**Affiliations:** Graduate Program in Neuroscience, University of Minnesota Twin Cities, Minneapolis, MN 55455, USA; Graduate Program in Molecular Pharmacology & Therapeutics, University of Minnesota Twin Cities, Minneapolis, MN 55455, USA; Department of Experimental and Clinical Pharmacology, University of Minnesota Twin Cities, Minneapolis, MN 55455, USA

**Keywords:** Alzheimer’s disease, human iPSC, APOE, HDL-mimetic peptide

## Abstract

APOE is the greatest genetic risk factor for late-onset Alzheimer’s disease (AD). In humans, APOE has three isoforms: APOE2 (E2), APOE3 (E3), and APOE4 (E4); E4 increases AD risk, while E3 is neutral and E2 decreases risk. In the brain, APOE is predominantly produced by astrocytes, where it binds lipids to form HDL-like particles, and plays a central role in lipid homeostasis, Aβ clearance, and neuroimmune modulation. Its lipidation state is critical for function, with E4 being poorly lipidated compared to E2 and E3, contributing to the pathogenic effects of E4 while also offering a potential therapeutic target. We have previously demonstrated that the HDL-mimetic peptide 4F increases APOE secretion and lipidation in wild-type mouse astrocytes and counteracts the inhibitory effects of Aβ42. Here, we assessed the ability of 4F to mitigate E4-associated dysfunction using primary astrocytes from humanized E3 and E4 knock-in mice and isogenic human iPSC-derived astrocytes and cerebral organoids. Results showed that 4F enhanced APOE secretion and lipidation in both cellular and organoid models in the absence or presence of aggregated Aβ42. Compared to E3 astrocytes, E4 astrocytes were prone to Aβ42-induced inhibition of APOE secretion and lipidation and increased accumulation of lipid droplets. 4F treatment ameliorated the inhibitory effects of Aβ42 and reduced lipid droplet accumulation. These findings support the therapeutic potential of HDL-mimetic peptides for E4-associated dysfunction in AD.

## Introduction

Alzheimer’s disease (AD) is the most common neurodegenerative disease and a leading cause of dementia worldwide (*Alzheimer’s Disease Facts and Figures.*, 2025; Mayeux & Stern, 2012). It is characterized pathologically by the formation of extracellular amyloid plaques that contain amyloid-β (Aβ), and intracellular tau neurofibrillary tangles (NFT) (Schachter & Davis, 2000) which eventually leads to severe neurodegeneration, brain atrophy, and cognitive dysfunction (Selkoe & Hardy, 2016). While 1-5% of AD cases are caused by genetic mutations in APP, PSEN1, and PSEN2, most AD cases are classified as sporadic, late-onset AD (LOAD) and influenced by multiple environmental and genetic risk factors (Andrade-Guerrero et al., 2023). Genome-wide association studies have identified many top risk genes for AD that play a role in lipid trafficking and metabolism, including APOE (Karch & Goate, 2015; Hollingworth et al., 2011; Holstege et al., 2021). The greatest genetic risk factor for developing LOAD is the APOE gene, which in humans has three alleles. APOE2 is protective against AD, APOE3 is neutral and the most common variant globally, and APOE4 increases AD risk, especially in homozygous carriers (Frisoni et al., 2022; Karch & Goate, 2015).

APOE is the most abundant apolipoprotein in the brain, where it is mainly produced and secreted by astrocytes and to a lesser extent microglia and neurons under stress conditions (Xu et al., 2006). It plays a central role in maintaining lipid homeostasis and other AD-related processes including Aβ aggregation and clearance, neuroimmune modulation, as well as mediating synaptic and cerebrovascular function (Kanekiyo et al., 2014; Koistinaho et al., 2004; Parhizkar & Holtzman, 2022). APOE exerts its function on lipid metabolism and transport by binding lipids (primarily phospholipids and cholesterol) to form high-density lipoprotein (HDL)-like particles in the interstitial and cerebrospinal fluid (Mahley, 1988). The degree of lipidation is important for APOE function and depends on its isoform. APOE2 and APOE3 are efficiently lipidation and facilitate toxic protein clearance, whereas APOE4 is poorly lipidated (Gong et al., 2002; Michikawa et al., 2000; Hanson et al., 2013; Hu et al., 2015).

Deficiency in lipid transport and proper redistribution has been linked to increased Aβ deposition and cholesterol in AD brains (Corraliza-Gomez et al., 2019; Xiong et al., 2007). Growing evidence from human studies, animal models, and cell models supports the beneficial role of HDLs in mitigating AD. HDLs exhibit robust lipid-binding and transport capabilities and have well-established protective effects on cardiovascular and brain function (Hottman et al., 2014). However, the therapeutic potential of native HDLs is limited due to their low bioavailability and inability to effectively cross the blood-brain barrier (BBB) (Leman et al., 2013). The development of small HDL mimetic peptides offers a promising alternative. Although they lack sequence homology to apolipoproteins, HDL mimetic peptides structurally mimic the class A amphipathic helices found in HDL-associated apolipoproteins such as APOA-I and APOE (Segrest et al., 1992; Leman et al., 2013). The most notable of these HDL mimetic peptides is the 18 amino acid peptide 4F, which contains 4 phenylalanine groups (Segrest et al., 1983; Datta et al., 2004). 4F has been clinically tested for cardiovascular diseases and atherosclerosis following successful studies in animal models (Navab et al., 2008). Similarly to HDLs, 4F possess anti-oxidative, anti-inflammatory, and antithrombotic properties (White et al., 2019). Several studies in mouse models as well as clinical trials have revealed the anti-inflammatory and anti-atherosclerosis properties of 4F (Navab et al., 2002; Lynch et al., 2003; Anantharamaiah et al., 2007; He et al., 2018). Recently, we have shown that 4F penetrates the BBB considerably more efficiently than the major circulating HDL protein APOA-I and enhances Aβ42 clearance from the brain (Swaminathan et al., 2020). Furthermore, 4F treatment reduces neuroinflammation, improves cognitive function, and mitigates both cerebral vascular and parenchymal amyloid pathology in mouse models of AD (Handattu et al., 2009; Zhong et al., 2025), supporting a beneficial role in the central nervous system.

However, the effects of 4F in the context of different APOE isoforms have not been investigated. We have previously shown that 4F increases APOE secretion and lipidation in primary mouse astrocytes and microglia, and primary human astrocytes (Chernick et al., 2018). In addition, 4F counteracts aggregated Aβ42-induced inhibition on APOE secretion and lipidation in astrocytes. Building upon our previous findings, the current study aims to evaluate the efficacy of 4F treatment in reversing APOE4-associated deficits using primary astrocytes from humanized APOE mice and isogenic human induced pluripotent stem cell (iPSC)-derived astrocytic and organoid models. Our results demonstrate that 4F enhances APOE secretion and lipidation in both primary and iPSC-derived models expressing human APOE, mitigates the inhibitory effect of Aβ42, and alleviates lipid droplet accumulation in APOE4 astrocytes.

## Methods

### Primary astrocyte culture

Homozygous humanized APOE KI mice expressing APOE3 (E3) or APOE4 (E4) were purchased from Jackson laboratory (Stock # 029018 and 027894). Primary glial cells were isolated from neonatal pups (postnatal day 1-3) as previously described (Chernick et al., 2018). Briefly, neonatal brains were dissected, and cortical tissues were isolated without the meninges. Cortical tissues were trypsinized into single cells and cultured on PDL-coated flasks for at least 2-3 weeks in Dulbecco’s Modified Eagle Medium (DMEM) supplemented with 10% fetal bovine serum, 16mM HEPES buffer, 0.1mM nonessential amino acids, 2 mM GlutaMAX, and 1X antibiotic/antimycotic. Purity of astrocyte cultures was determined by immunofluorescence. All animal procedures were approved by the Institutional Animal Care and Use Committee (IACUC protocol # 2207-40221A) at the University of Minnesota.

### iPSC maintenance and differentiation

Isogenic APOE3 and APOE4 iPSC’s derived from a homozygous *APOE3* individual were generously provided by Dr. Matthew Blurton Jones. iPSCs were maintained on vitronectin coated plates, cultured in E8 flex media and passaged every 3-4 days using citrate buffer. iPSCs were routinely checked for mycoplasma. To initiate differentiation into neural progenitor cells (NPCs), iPSC colonies were enzymatically dissociated using accutase and seeded onto Matrigel-coated 6 well plates. iPSCs were fed daily with Neural induction medium (Stemcell Technologies) supplemented with LDN 193189 (Cayman Chemicals) and SB 431542 (Selleckchem) for 7-10 days. NPCs were maintained as neural rosettes in neural maintenance medium (DMEMF12 medium with N2 and B27 supplements) with bFGF until further differentiation.

Differentiation into astrocytes was accomplished following an established protocol (TCW et al., 2017). Briefly, NPCs were plated at low density (15k/cm2) onto Matrigel-coated plates. Cells were fed every 2-3 days with astrocyte media (Sciencell) supplemented with Astrocytes Growth Serum, Fetal Bovine Serum, and Pennicilin/Streptomycin, and passaged with accutase every 10-14 days, or when confluent. Mature astrocytes were plated onto Matrigel coated dishes for experiments at day 40-60. Differentiated cells were validated by staining for the NPC markers SOX2/Nestin and mature astrocyte markers GFAP and S100β.

Cerebral organoids were generated using previously published methodologies (Lindborg et al., 2016). Briefly, three 6-well plates of iPSC’s were dissociated and placed in a Cell-Mate 3D cocoon consisting of hyaluronic acid and chitosan to form an embedded matrix. Matrices were cultured in G-Rex flasks (Wilson Wolf) with E8 medium. Medium was added every 3-4 days for the first 2-3 weeks, or until organoids first started to emerge from the matrices. Half medium changes were performed thereafter. Organoids were used for experiments at 9-12 weeks.

### Treatments with 4F and Αβ42

The peptide 4F (Ac-DWFKAFYDKVAEKFKEAF-NH2) (4F: peptide purity=95%, peptide content=72%) was custom synthesized and purchased from American Peptide Company Inc (Sunnyvale, CA). 4F was prepared and used for treatment as previously described (Chernick et al., 2018). Briefly, astrocytes were treated with 5 μM 4F or PBS as a control for 24 hours in minimal volume of serum-free Opti-MEM, supplemented with antibiotics and 0.1% BSA. Synthetic Aβ42 was purchased from Advanced Automated Peptide Protein Technologies (Louisville, KY). Αβ42 aggregates were prepared as previously described (Chernick et al., 2018; Stine et al., 2011). Briefly, 0.05 mg of peptide were dissolved in HFIP for at least 30 min, and then aliquoted and allowed to dry overnight. Dried peptides were then placed in speed vacuum for at least 1 hour to create peptide films. Peptide films were dissolved to 5 mM in DMSO, sonicated in a water bath, and then mixed with phenol free DMEM to a final concentration of 100 μM. The solution was vortexed and incubated at 37°C overnight to form aggregated Αβ42. Cells were treated with Αβ42 aggregates for 24 hours at 5 μM final concentration.

### Immunofluorescence and immunohistochemistry

Cells were plated in 48-well plates and allowed to adhere overnight. Cells were fixed with 4% PFA, permeabilized for 10 minutes in 0.5% TritonX-100 and blocked for 1 hour in PBS with 0.05% tween (PBST) + 5% serum. Cells were incubated overnight in blocking solution with the following primary antibodies: SOX2 (Cell Signaling 3579), Nestin (Invitrogen MA1-110), GFAP (Dako Z0334), S100β (Millipore S2532), and MAP2 (Invitrogen PA1-10005). Organoids were fixed overnight, washed, and stored in 70% ethanol for paraffin embedding. Immunohistochemistry for MAP2 and GFAP was performed on paraffin-embedded organoid sections using standard protocols at the University of Minnesota Comparative Pathology Shared Resource.

### BODIPY staining

Astrocytes were plated onto Matrigel-coated 48 wells and allowed to adhere overnight. Astrocytes were treated with 30 μM oleic acid or 0.1% fatty acid-free BSA as a control for 24 hours in the presence or absence of 4F. Astrocytes were fixed with 4% PFA, permeabilized for 10 minutes in 0.5% TritonX-100 and stained with BODIPY 493/503 (a fluorescent dye staining neutral lipids; Thermofisher) for 1 hour. Cells were cover slipped with DAPI and imaged using a Keyence microscope.

### Western blotting and non-denaturing gel electrophereses

Conditioned media was collected with the addition of protease inhibitors AEBSF and aprotinin. Cells were washed with PBS and harvest in RIPA buffer supplemented with protease and phosphatase inhibitors. Cell lysates were extracted by sonication, at 15% amplitude 3x 10 seconds each. Lysates were centrifuged at 12,000G at 4°C and supernatant was collected. Equal volume of media or of RIPA lysates supplemented with 5% BME were boiled, loaded onto a 12% tris-glycine gel, and separated via SDS-PAGE. Gels were transferred onto PVDF membranes for 1.5-2 hours. Membranes were blocked with 5% milk in PBST for 1 hour. Primary antibodies were incubated overnight at 4°C and detection was carried out with HRP-conjugated secondary antibodies and developed using chemiluminescence (Biorad). Membranes were scanned using the iBright imager (Thermofisher). For non-denaturing native gel electrophoresis (NDGGE), fresh (not frozen) media was mixed with native sample buffer without SDS or BME. Media samples were loaded without boiling into a 4-20% gradient tris-glycine gel and run for 5 hours. Membranes were then transferred, blocked and incubated with APOE antibody (Millipore 178479) as described above. Western blot results were quantified using Image J software (National Institutes of Health, Bethesda, MA, USA). Secreted APOE was determined as a ratio of APOE present in media: APOE present in RIPA fraction. HMW particles (10-17 nm) were considered lipidated APOE, while LMW (∼8.2 nm) particles were considered poorly lipidated. Lipidated APOE was determined as a ratio of HMW/ total lipidation. Data are expressed as relative percent, with vehicle set to 100%.

### Statistical analysis

Data are expressed as Mean ± standard error (SEM) from at least 3 independent experiments with treatments performed in triplicates in each independent experiment, unless otherwise indicated. Data was analyzed by Student’s t-test/ Tukey’s one-way ANOVA (when normally distributed), or Mann-Whitney/ Wilcoxon rank sum test and Kruskal Wallis test (when not normally distributed). The p value < 0.05 was considered statistically significant. Statistical tests were carried out using Graphpad Prism.

## Results

### Reduced APOE secretion and lipidation in APOE4 primary astrocytes

APOE4 has been shown to be less lipidated compared to other APOE genotypes (Gong et al., 2002; Michikawa et al., 2000; Hanson et al., 2013; Hu et al., 2015). To determine whether the APOE4-associated lipidation deficit remains true in our cellular models, we cultured primary astrocytes from humanized APOE4 and APOE3 knock-in (KI) mice following our established protocol. We found that APOE4 primary astrocytes displayed a modest decrease in APOE secretion (**Figure 1A**) but a significant reduction (approximately 50%) in APOE lipidation compared to APOE3 (**Figure 1B**).

**Figure 1.**
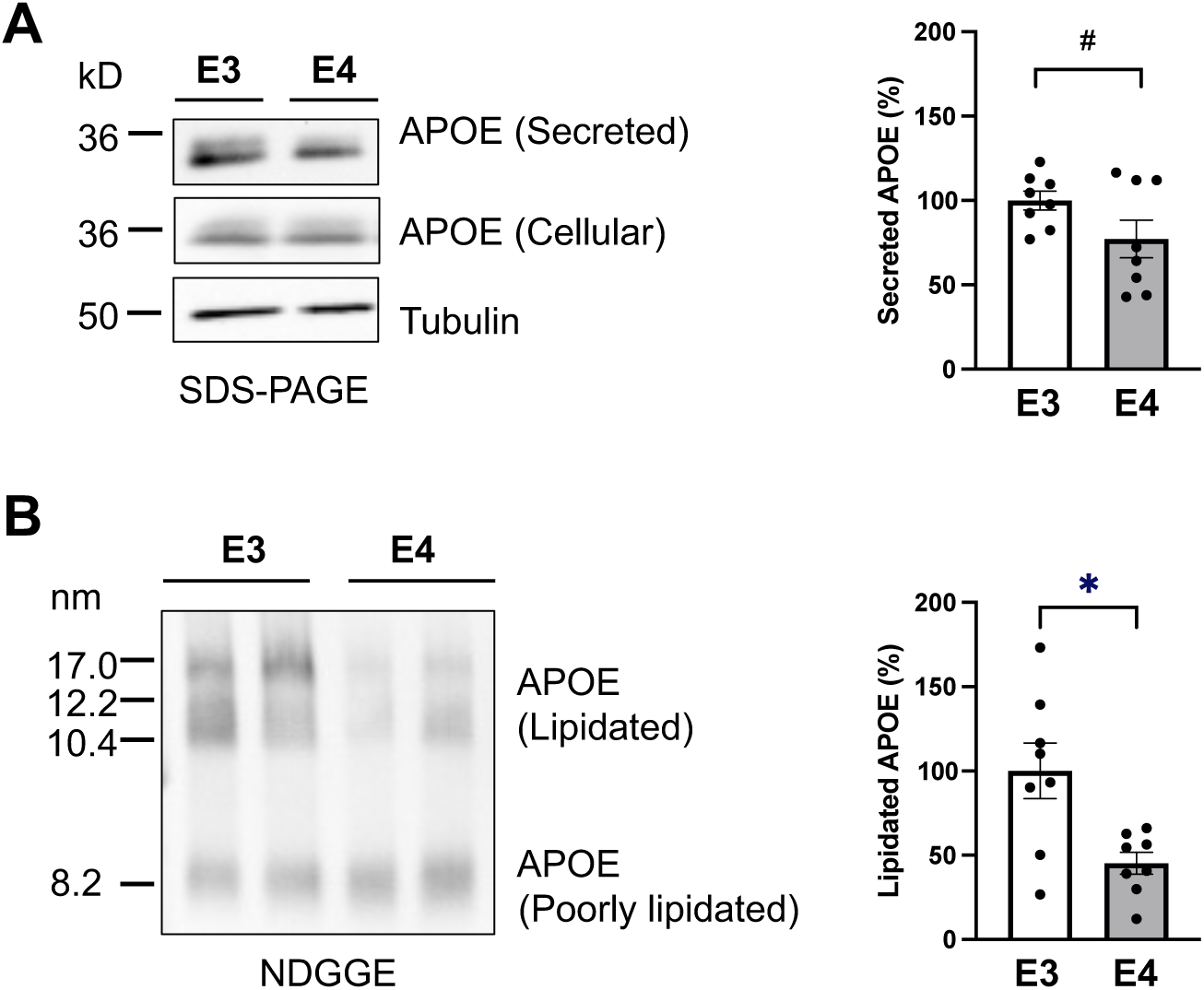
APOE4 primary astrocytes exhibit decreased APOE secretion and lipidation. Primary astrocytes from humanized APOE3 or APOE4 mice were incubated in serum-free media for 24 hours. SDS-PAGE and NDGGE were performed to determine the relative secretion and lipidation state of APOE. (**A**) Representative image and quantification from SDS-PAGE performed on media and cell lysates from primary astrocytes. (**B**) Representative image and quantification from NDGGE performed on fresh media from primary astrocytes. Tubulin was used as a loading control. Data was normalized to APOE3 and expressed as a percent. Data represents Mean ± SEM of 7-8 replicates from 3 independent experiments. Welsch’s t-test, *p < 0.05, **p < 0.01, ***p < 0.001.

### 4F Treatment rescues lipidation deficits in primary astrocytes and enhances APOE secretion and lipidation in primary and iAstrocytes

To investigate the impact of 4F on APOE secretion and lipidation, APOE3 and APOE4 primary astrocytes were subjected to 4F treatment for 24 hours. 4F treatment significantly enhanced APOE secretion (**Figure 2A**) and lipidation (**Figure 2B**) in both APOE3 and APOE4 astrocytes. In parallel, we differentiated isogenic human iPSCs homozygous for APOE3 or APOE4 into astrocytes (iAstrocytes), following the published protocols (TCW et al., 2017). iPSCs were first differentiated into neural progenitor cells (NPCs) expressing SOX2 and Nestin, then further matured into astrocytes expressing mature markers GFAP and S100b (**Figure S1A**). Interestingly, baseline APOE secretion and lipidation levels were comparable between the two genotypes in the iAstrocytes (**Figure 3A,B,C,D**), indicating that the lipidation deficit seen in primary APOE4 astrocytes may depend on specific model systems and cell lines. However, similar to our findings in primary astrocytes, 4F treatment markedly increased APOE secretion (**Figures 3A,E**) and lipidation (**Figure 3A,F**) in APOE3 and APOE4 iAstrocytes. Overall, the results showed that 4F promoted APOE secretion and lipidation in both APOE3 and APOE4 primary astrocytes and iAstrocytes, suggesting that 4F may enhance APOE3 function in addition to rescuing APOE4 deficits.

**Figure 2.**
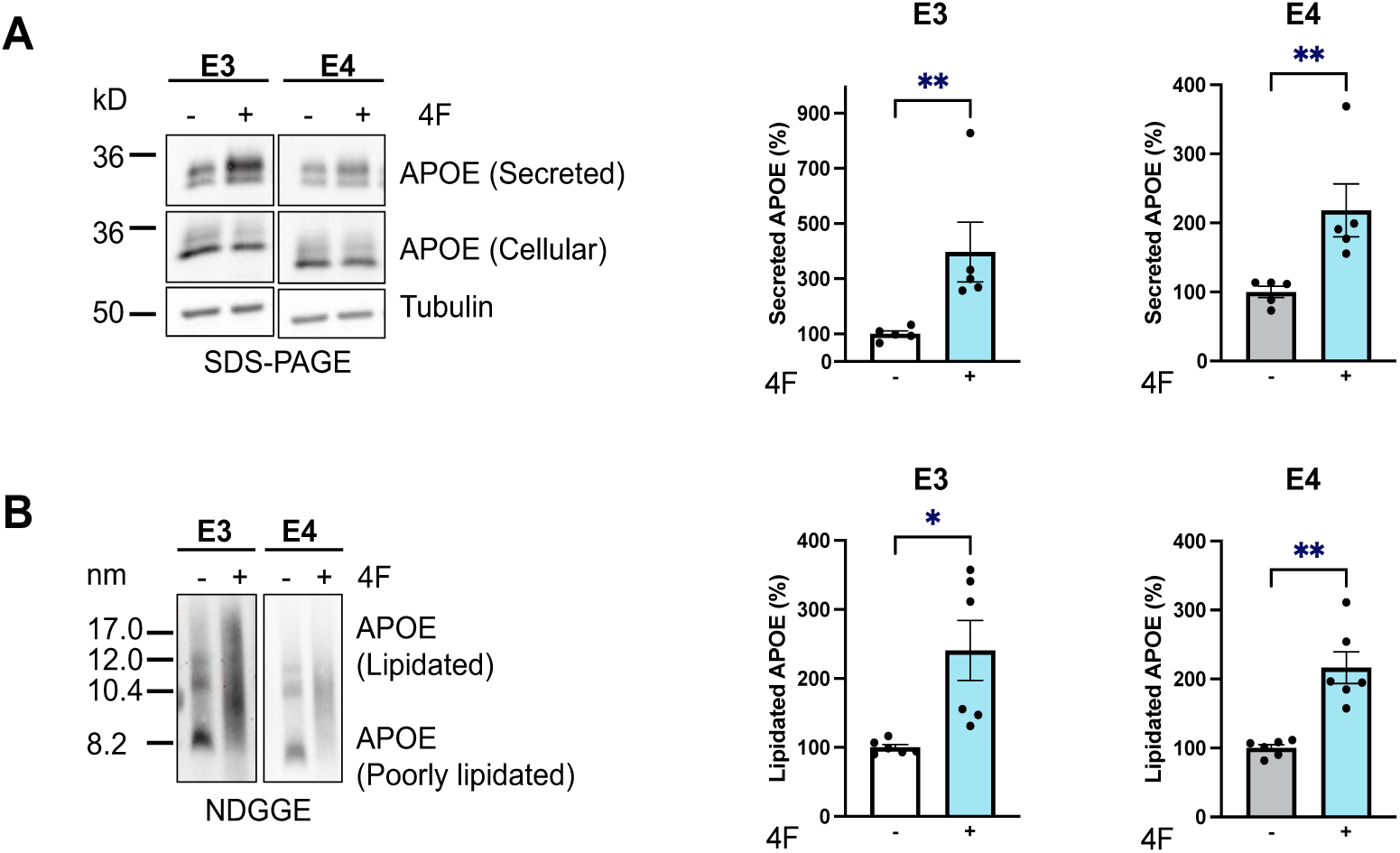
4F Treatment rescues lipidation deficits in primary astrocytes and enhances APOE secretion and lipidation in primary astrocytes. (A) Representative image and quantification from SDS-PAGE performed on media and cell lysates from primary astrocytes. (B) Representative image and quantification from NDGGE performed on fresh media from primary astrocytes. Tubulin was used as a loading control. Data was normalized to vehicle for each genotype and expressed as a percent of vehicle treatment. Data represents Mean ± SEM of 7-12 replicates from 3 independent experiments. Welsch’s t-test, *p < 0.05, **p < 0.01, ***p < 0.001.

**Figure 3.**
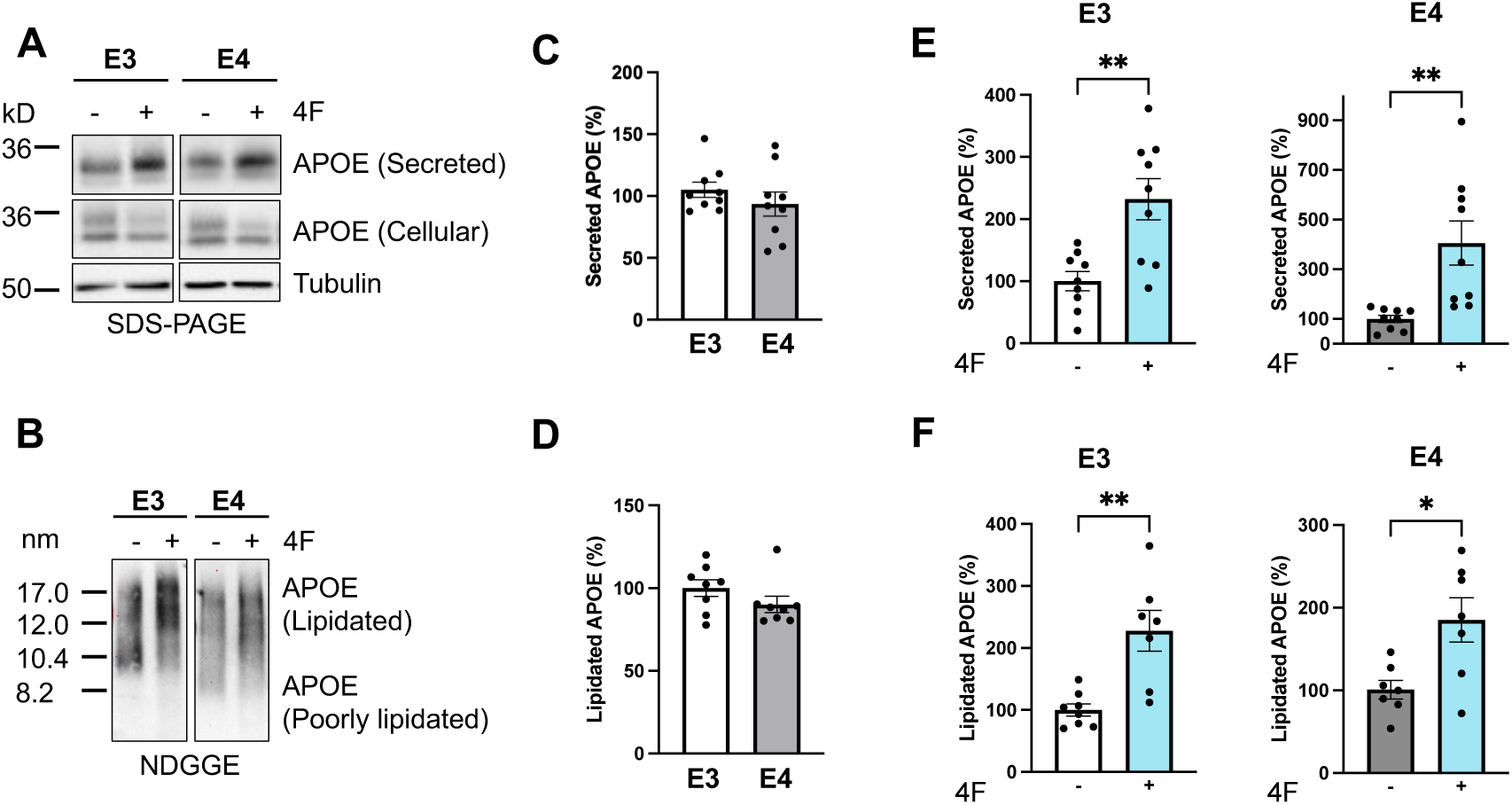
4F treatment enhances APOE secretion and lipidation in iAstrocytes. (A) Representative image from SDS-PAGE performed on media and cell lysates from iAstrocytes. (**B**) Representative image from NDGGE performed on fresh media from primary astrocytes. (**C,D**) Quantification of secreted and lipidated APOE from iAstrocytes. Data was normalized to APOE3 and expressed as a percent. (**E,F**) Quantification of secreted and lipidated APOE from iAstrocytes treated with 4F. Tubulin was used as a loading control. Data was normalized to vehicle for each genotype and expressed as a percent of vehicle treatment. Data represents Mean ± SEM of 7-12 replicates from 3 independent experiments. Welsch’s t-test, *p < 0.05, **p < 0.01, ***p < 0.001.

### Aβ42 inhibits APOE secretion/lipidation and 4F treatment counteracts Aβ42-induced inhibitory effects in APOE4 astrocytes

Previous studies have shown that aggregated Aβ42 inhibits APOE secretion and lipidation, which is counteracted by a co-treatment with 4F in wild-type mouse primary astrocytes (Chernick et al., 2018). To determine the effects of Αβ42 and 4F on APOE secretion and lipidation across different APOE isoforms in humanized APOE models, primary astrocytes and iAstrocytes were treated with aggregated Αβ42 without or with 4F for 24 hours. The data showed that treatment with aggregated Aβ42 decreased APOE secretion (**Figure 4A**) and lipidation (**Figure 4B**) in APOE4 primary astrocytes, as in primary mouse astrocytes expressing endogenous mouse APOE (Chernick et al., 2018). APOE3 astrocytes however were more resistant to the impact of Aβ42 and did not exhibit significant changes in APOE secretion and lipidation (**Figure 4A,B**). Notably, the Aβ42-induced reduction in APOE secretion and lipidation in APOE4 primary astrocytes was observed despite their already low baseline levels (**Figure 1A, B**; **Figure 4A,B**), indicating an additive inhibitory effect of Aβ42 on APOE secretion and lipidation in APOE4 astrocytes. Remarkably, 4F treatment increased APOE secretion (**Figure 4A**) and lipidation (**Figure 4B**) in both APOE3 and APOE4 primary astrocytes in the presence of aggregated Aβ42. In particular, 4F treatment counteracted the inhibitory effects of Aβ42 and increased APOE4 secretion and lipidation in E4 primary astrocytes to the similar levels as in E3 primary astrocytes (**Figure 4A,B**).

**Figure 4.**
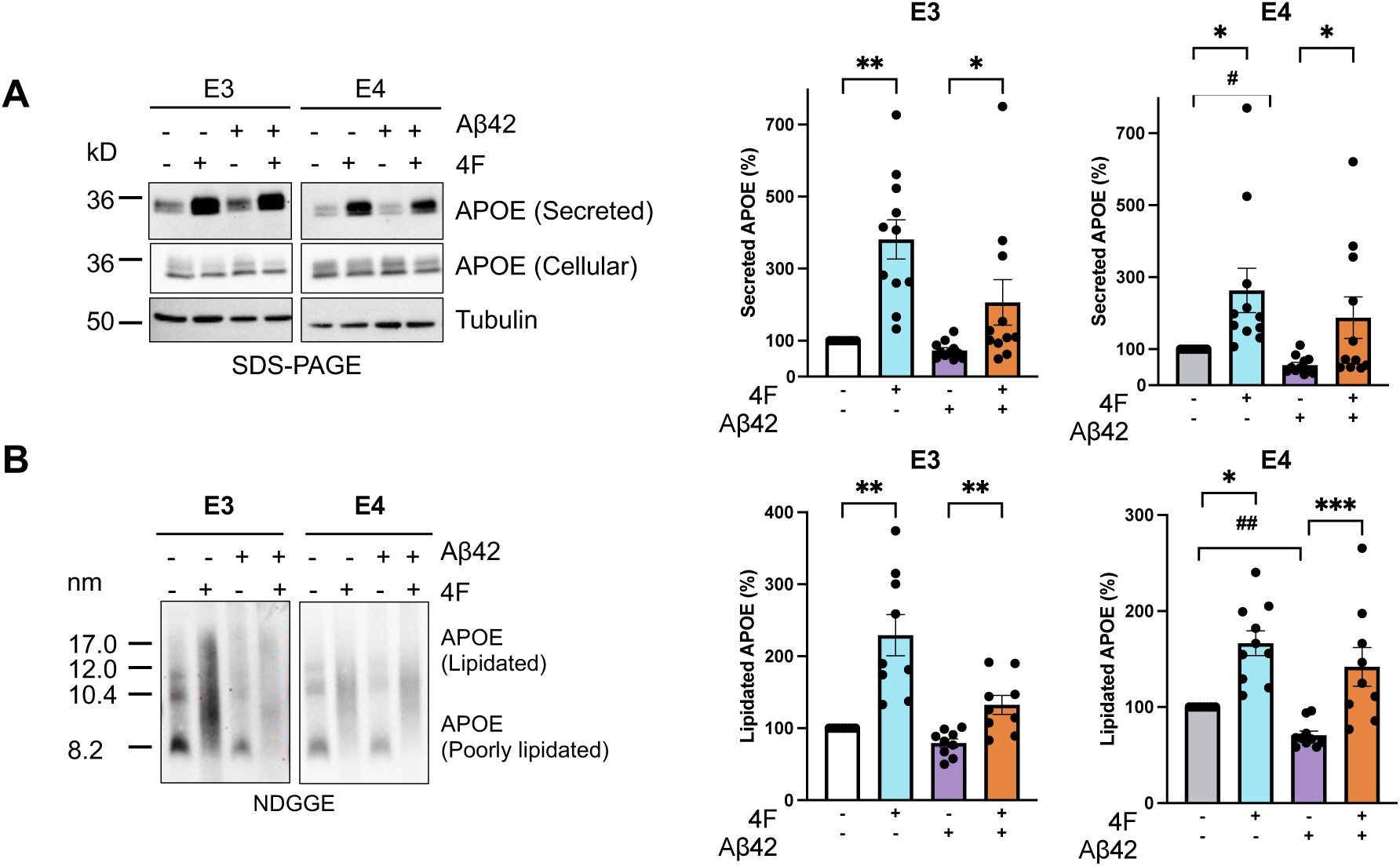
4F rescues the inhibitory effect of Αβ42 on APOE secretion and lipidation in primary APOE4 astrocytes. (**A**) Representative image and quantification from SDS-PAGE performed on media and cell lysates in primary astrocytes. (**B**) Representative image and quantification from NDGGE performed on media in primary astrocytes. Tubulin was used as a loading control. Data was normalized to vehicle for each genotype and expressed as a percent of vehicle treatment. Data represents Mean ± SEM of 7-9 replicates from 3 independent experiments. One-way ANOVA with Tukey’s post-hoc correction, #p = .052, ##p= .06 * p < .05, ** p < .01, *** p < .001

In iAstrocytes, although APOE4 astrocytes did not exhibit a deficit in APOE secretion and lipidation compared to APOE3 astrocytes under basal conditions (**Figure 2A**), upon Aβ42 treatment, APOE secretion was significantly decreased in APOE4 iAstrocytes whereas there was no change in APOE3 iAstrocytes (**Figure 5A**). Interestingly, Aβ42 treatment did not seem to affect APOE lipidation in APOE3 or APOE4 iAstrocytes, although the effects might be masked by relatively low basal levels of lipidated APOE in iAstrocytes (**Figure 5B**). Importantly, 4F treatment promoted APOE lipidation in the presence of aggregated Aβ42 to similar extents in APOE3 and APOE4 iAstrocytes (**Figure 5B**).

**Figure 5.**
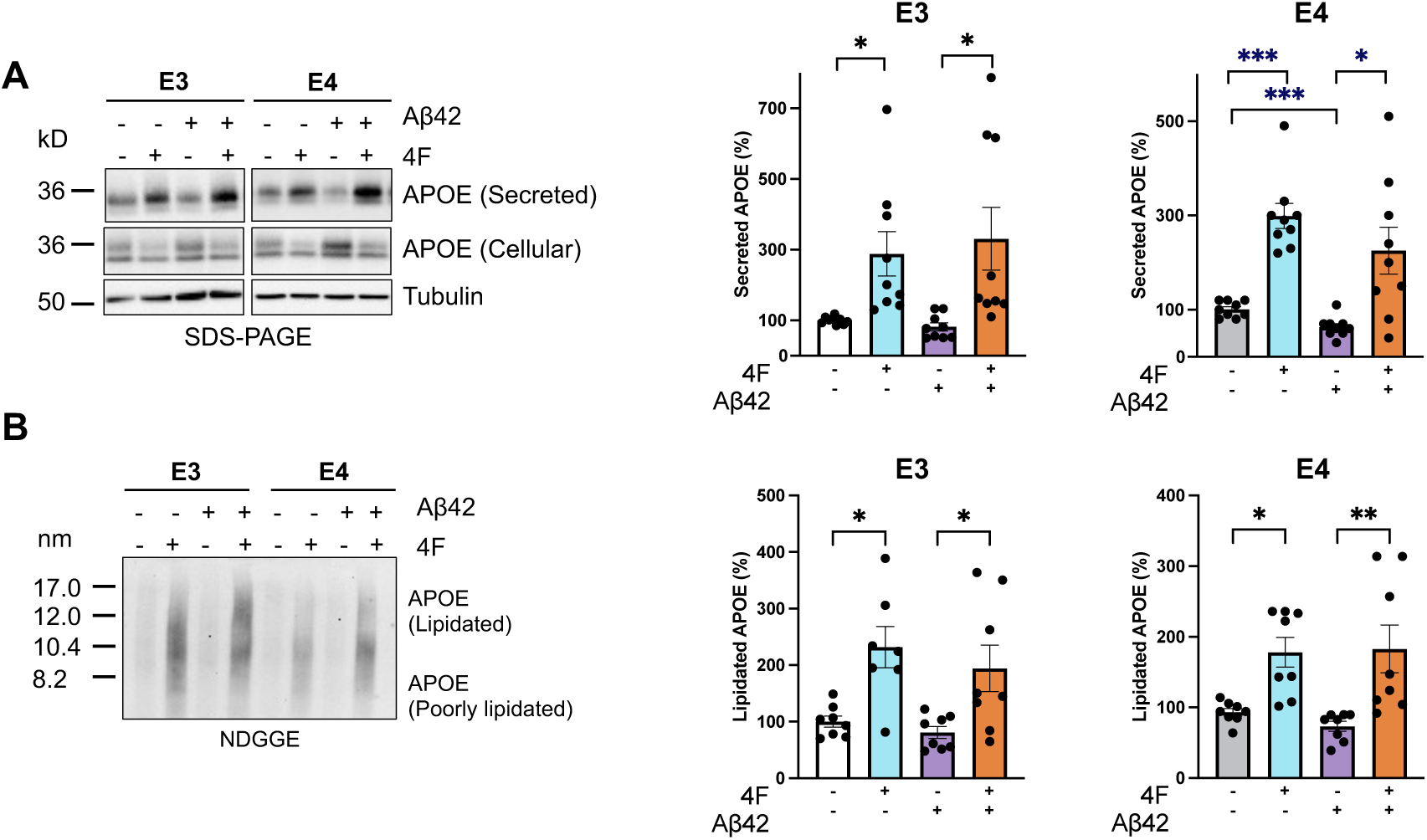
4F rescues the inhibitory effect of Αβ42 on APOE secretion and lipidation in APOE4 iAstrocytes. (**A**) Representative image and quantification from SDS-PAGE performed on media and cell lysates in iAstrocytes. (**B**) Representative image and quantification from NDGGE performed on media in iAstrocytes. Tubulin was used as a loading control. Data was normalized to vehicle for each genotype and expressed as a percent of vehicle treatment. Data represents Mean ± SEM of 7-9 replicates from 3 independent experiments. One-way ANOVA with Tukey’s post-hoc correction, #p = .052, * p < .05, ** p < .01, *** p < .001

Taken together, compared to APOE3 astrocytes, APOE4 astrocytes were prone to the inhibition of Αβ42 on APOE secretion and lipidation, and 4F treatment counteracted Aβ42-induced inhibitory effects in both APOE3 and APOE4 primary and iAstrocytes. These findings highlight the potential resilience that 4F treatment may provide against the pathological inhibition imposed by amyloid aggregation on APOE function, particularly in APOE4 astrocytes, which are more susceptible to Aβ42.

### 4F mitigates APOE4-induced cholesterol/lipid metabolism dysfunction in iAstrocytes

Excess neutral lipids including triglycerides and cholesterol esters are stored within intracellular organelles as lipid droplets (LDs). LDs accumulate under conditions of stress, aging and in AD brains (Haney et al., 2024; Marschallinger et al., 2020; Smolič et al., 2021). In addition, previous studies have demonstrated that APOE4 astrocytes exhibit aberrant lipid trafficking and accumulate cholesterol and unsaturated triglycerides in the form of lipid droplets (Farmer et al., 2020; Qi et al., 2021; Windham et al., 2024). APOE4 astrocytes also reduce neuronal sequestering of LDs, contributing to synaptic dysfunction (Ioannou et al., 2019; Qi et al., 2021).

To investigate whether 4F could attenuate lipid droplet accumulation, APOE3 and APOE4 iAstrocytes were subjected to BODYPI staining and imaging analyses. Under basal conditions, APOE4 astrocytes accumulated significantly more lipid droplets than APOE3 astrocytes (**Figure 6A,B**). Treatment with 4F at the basal condition did not significantly alter lipid droplets. However, upon challenged with oleic acid treatment, which promotes lipid droplet formation (Listenberger et al., 2016), APOE4 astrocytes exhibited a further increase in lipid droplet accumulation compared to APOE3 astrocytes (**Figure 6A,B**). Remarkably, 4F treatment significantly reduced lipid droplet accumulation in both APOE4 and APOE3 astrocytes under oleic acid challenge (**Figure 6A,B**). Taken together, these findings suggest that 4F reduces intracellular lipid accumulation while enhancing cholesterol/lipid efflux for APOE lipidation in iAstrocytes.

**Figure 6.**
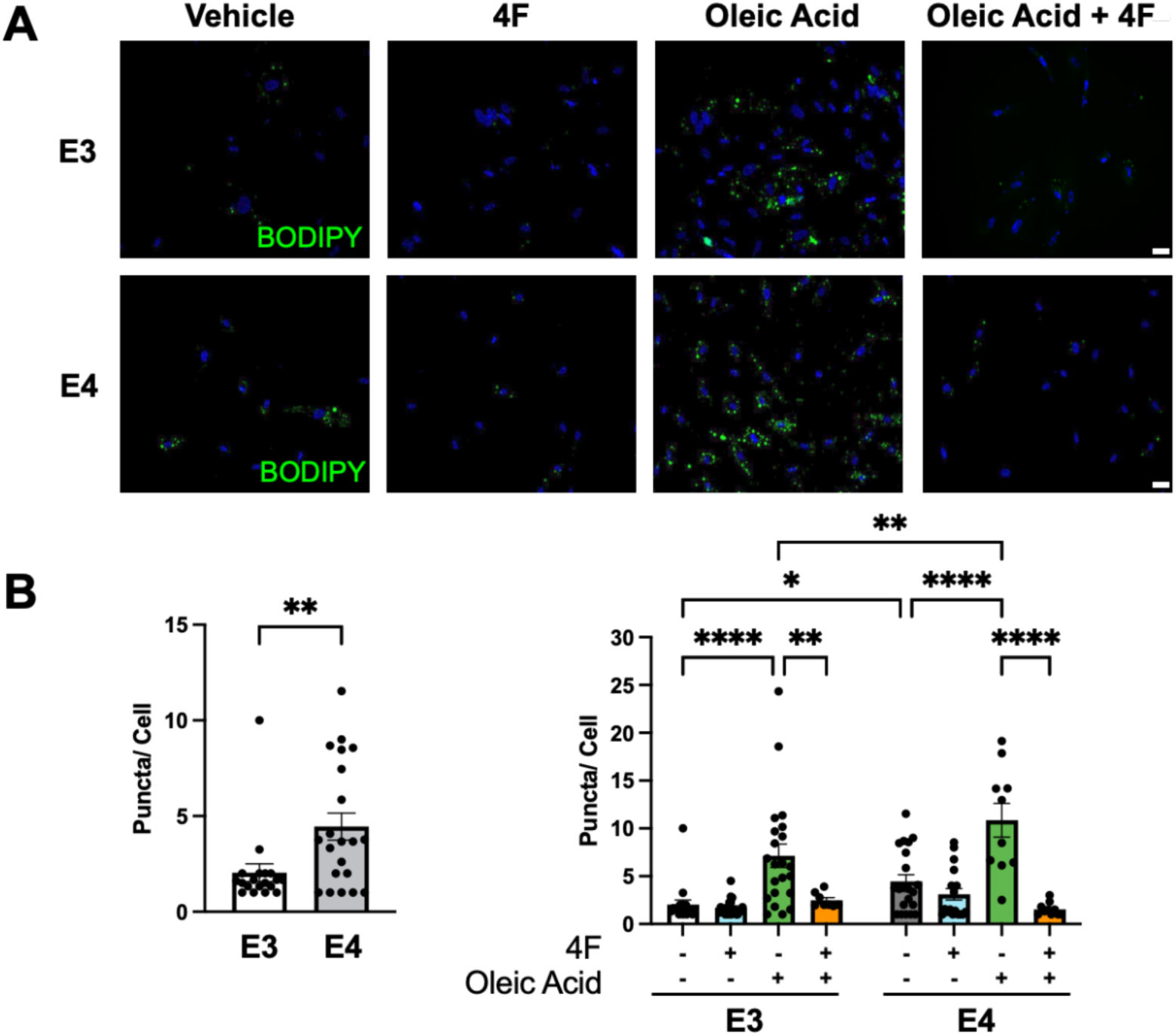
4F mitigates APOE4-related cholesterol/lipid metabolism dysfunction in iAstrocytes. APOE3 and APOE4 iPSC-derived astrocytes were treated with 30 μM oleic acid with or without 5 μM 4F for 24 hours and stained with BODIPY to measure lipid droplets. Number of puncta were quantified and normalized to DAPI. Data represents average number of puncta/cell per field of view from 3 culture wells from 1 representative experiment. Scale bar, 50 μm. (**A**) Representative images and (**B**) quantification of APOE3 or APOE4 iAstrocytes treated with vehicle or 4F +/- oleic acid. Two-way ANOVA with Tukey’s post-hoc correction. * p < .05, ** p < .01, **** p <.0001

### 4F enhances secretion and lipidation in iPSC-derived cerebral organoids in the absence or presence of Αβ42

To explore the effects of 4F treatment in a complex, multicellular system, APOE3 and APOE4 iPSC-derived cerebral organoids, which contain diverse cell types, including neurons and astrocytes, were generated (**Figure S1 B**), following an established differentiation protocol (Lindborg et al., 2016). At 2-3 months, the organoids were treated with 4F for 24 hours, followed by assessment of APOE secretion and lipidation. The results showed that 4F treatment promoted APOE secretion and lipidation in both APOE3 and APOE4 cerebral organoids (**Figures 7A,B**). To further assess the robustness of this effect, organoids were treated with 4F in the presence of Aβ42. Consistent with the observed results in astrocytes cultures, 4F increased APOE secretion and lipidation in the presence of Αβ42 in cerebral organoids, and these effects were particular evident in APOE4 organoids (**Figure 7A,B**). These findings demonstrate that 4F treatment can exert its effects on APOE secretion and lipidation in complex 3D organoids as well as in 2D cellular models.

**Figure 7.**
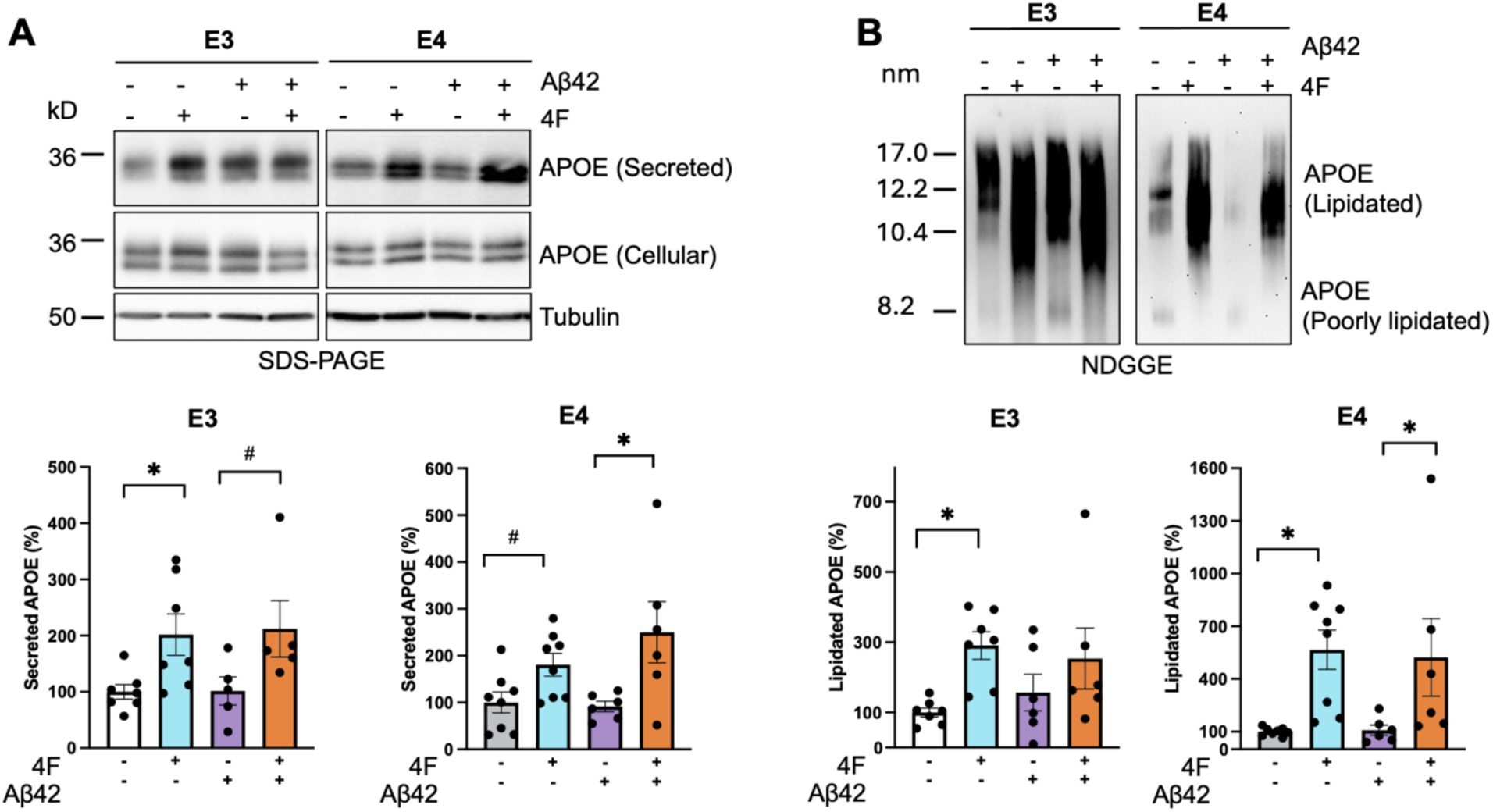
4F enhances secretion and lipidation in the presence and absence of Aβ42 in cerebral organoids. iPSCs were used to generate cerebral organoids. APOE3 and APOE4 cerebral organoids were treated with 5 μM 4F with or without 5 μM aggregated Aβ42 peptide for 24 hours. (**A**) Media and cell lysates were subjected to SDS-PAGE and immunoblot analysis. (**B**) Fresh media was analyzed for APOE lipidation using non-denaturing gradient gel electrophoresis (NDGGE). Data was normalized to vehicle for each genotype and expressed as a percent of vehicle treatment, Data represents Mean ± SEM of 7-8 replicates from 2 independent experiments. One-way ANOVA with Tukey’s post-hoc correction, # p = .06,* p < .05, **p < .01, ***p < .001.

## Discussion

This study aimed to determine the effects of the HDL mimetic peptide 4F on APOE secretion and lipidation in various APOE isoform-specific cellular models, including primary astrocytes from humanized APOE mice, isogenic human APOE iPSC-derived astrocytes and cerebral organoids. 4F enhanced APOE secretion and lipidation in both APOE4 and APOE3 primary and iAstrocytes, and in cerebral organoids as well. Exposure to Aβ42 exacerbates APOE secretion and lipidation deficits in APOE4 astrocytes. Importantly, 4F treatment rescued the inhibitory effect of Aβ42 on APOE secretion and lipidation in APOE4 astrocytes, and reduced lipid droplet accumulation. The results provide significant insights into isoform-dependent APOE secretion and lipidation as well as the differential effects of 4F treatment across these models.

Secretion and lipidation state of APOE is essential for its functions in lipid transport, synaptic maintenance, and neuroprotection (Hubin et al., 2019; Lanfranco et al., 2021) and is highly correlated with AD severity (Merched et al., 2000). APOE4 astrocytes exhibit reduced lipidation efficiency, impaired cholesterol efflux and decreased amyloid clearance (TCW et al., 2022). Numerous studies have demonstrated APOE4-associated lipid metabolism dysfunction and aberrant cholesterol homeostasis (De Leeuw et al., 2022; Jeong et al., 2019; Lindner et al., 2022; Rawat et al., 2019), underscoring the critical role of APOE secretion and lipidation in maintaining brain lipid homeostasis. In this study, we observed a clear deficit in APOE secretion and lipidation in primary APOE4 astrocytes derived from humanized APOE mice, consistent with previous reports of APOE4-associated dysfunction. In contrast, no significant baseline differences in secretion or lipidation were observed between APOE3 and APOE4 iAstrocytes, highlighting potential context-dependent effects of APOE genotype. Importantly, 4F treatment significantly increased APOE lipidation in both primary and iAstrocytes, as in wild-type mouse and human primary astrocytes reported previously (Chernick et al., 2018). These model-specific differences may reflect variability in cholesterol metabolism across model systems. Humanized APOE knock-in primary astrocytes express human APOE on a murine background, which may influence lipid metabolism differently compared to iAstrocytes. This supports prior findings that cellular contexts and model systems influence APOE-dependent lipid pathways (TCW et al., 2022) and underscores the importance of using diverse models to understand APOE4-related dysfunction.

One of the central mechanisms of AD pathogenesis is the interaction between APOE and Aβ, where APOE is known to influence both aggregation and clearance of Aβ (Bilousova et al., 2019; LaDu et al., 1994; Strittmatter et al., 1993; Tai et al., 2013, 2014). Aβ impairs APOE secretion and lipidation, disrupting lipid metabolism, neuroinflammation, amyloid clearance, synaptic function, and cerebrovascular health (Azizidoost et al., 2022). Although baseline APOE secretion and lipidation was not impaired in APOE4 iAstrocytes, exposure to aggregated Aβ42 induced a significant inhibition of APOE secretion, indicating a heightened vulnerability to pathological stressors in these cells. Similarly, APOE4 primary astrocytes displayed increased vulnerability to Aβ42-induced inhibition, compared to APOE3 primary astrocytes in which Aβ42 had no significant effects on APOE secretion or lipidation. Remarkably, 4F treatment counteracted Aβ42-induced inhibition in both primary and iAstrocytes, demonstrating a protective effect against Aβ-induced dysfunction. Notably, 4F also increased secretion and lipidation in APOE3 astrocytes exposed to Aβ42. This suggests that 4F can enhance APOE function under pathological stress across APOE genotypes, highlighting its therapeutic potential beyond APOE4 carriers. This broad effect is particularly promising for sporadic, late-onset Alzheimer’s disease (LOAD), where multiple genetic and environmental factors contribute to disease risk and lipid dysregulation alongside APOE4. Our data suggest that 4F could potentially provide benefits across diverse patient populations by improving APOE lipidation and function irrespective of APOE genotype.

Our findings align with prior studies demonstrating that 4F enhances APOE secretion and lipidation in primary mouse glia and in human astrocytes (Chernick et al., 2018). A recent study also showed that 4F enhances APOE lipidation and rescues cholesterol accumulation in NPC1-inhibited fibroblasts and iAstrocytes (Di Biase et al., 2025). Notably, recombinant APOE4 was less effective at normalizing cholesterol and endogenous APP levels in fibroblasts than APOE3, which was rescued with 4F treatment (Di Biase et al., 2025). Our study extends these findings by demonstrating that 4F similarly enhances APOE secretion and lipidation in isogenic APOE-isoform specific iAstrocytes and reverses APOE4-related deficits induced by extracellular Aβ42.

A plausible mechanism for the beneficial effects of 4F involves ATP-binding cassette transporter A1 (ABCA1), a key transporter mediating cholesterol/lipid efflux and generation of HDL-like lipoproteins (Boehm-Cagan et al., 2016; Islam et al., 2018; Jacobo-Albavera et al., 2021; Larrede et al., 2009). Prior studies have shown that HDL mimetic peptides interact with and stabilizes ABCA1, facilitating cholesterol/lipid efflux and APOE lipidation (Arakawa et al., 2004; Valencia-Olvera et al., 2023). ABCA1 is essential for 4F action, as ABCA1 knockout abolishes 4F-induced increases in APOE secretion and lipidation (Chernick et al., 2018). Moreover, ABCA1 deletion exacerbates Aβ pathology in AD mouse models (Wahrle et al., 2005, 2008). Interestingly, prior studies reported no changes in ABCA1 protein levels following 4F treatment in primary astrocytes (Chernick et al., 2018), suggesting that 4F may act on ABCA1 through primarily post-translational mechanisms to enhance cholesterol/lipid efflux and APOE lipidation.

Lipid droplets (LDs) are intracellular organelles that store neutral lipids such as triglycerides and cholesterol esters. Under normal physiological conditions, lipid droplets facilitate temporary storage of triglycerides and cholesterol esters until they undergo lipolysis or lipophagy or their contents are exported via HDL-like particles (Olzmann & Carvalho, 2019). APOE4 astrocytes in AD exhibit impaired Aβ uptake, increased cholesterol accumulation and enhanced LD formation along with decreased cholesterol efflux capacity, contributing to lipid metabolism dysfunction. In addition, APOE4 impairs synaptic support of astrocytes to neurons (Gong et al., 2002; Michikawa et al., 2000; TCW et al., 2022). Consistent with previous findings (Sienski et al., 2021), APOE4 iAstrocytes showed more LD accumulation than APOE3 iAstrocytes at baseline and following fatty acid challenge in the present study. Importantly, 4F treatment significantly attenuated LD accumulation in both APOE3 and APOE4 iAstrocytes under fatty acid challenge, suggesting that 4F promotes intracellular lipid homeostasis, particularly in APOE4 astrocytes. Our data corroborate and extend increased LD accumulation as an APOE4-dependent phenotype that is exacerbated by fatty acid challenge, which can be ameliorated by 4F treatment, supporting the potential of 4F to restore APOE4-driven lipid dysregulation.

Targeting APOE lipidation is an emerging therapeutic approach for AD in preclinical and cellular models (Patil & Kuehn, 2024). Strategies that enhance cholesterol/lipid efflux, including APOA-I and other APO/HDL mimetic peptides, have shown benefits in AD models (Chernick et al., 2020). CS-6253, a peptide derived from APOE’s C-terminus, has been reported to activate ABCA1 and promote APOE lipidation and reduces amyloid burden in vivo (Boehm-Cagan et al., 2016; Valencia-Olvera et al., 2023). Liver X receptor (LXR)/retinoid X receptor (RXR) agonists, which upregulate ABCA1 expression (Larrede et al., 2009), also enhance lipidation and mitigate amyloid pathology and neuroinflammation (Koldamova et al., 2005; Zelcer et al., 2007; Donkin et al., 2010; Cramer et al., 2012). Notably, HDL mimetic peptides including 4F have consistently demonstrated cognitive and neuroprotective benefits in animal models of AD and brain injury (Handattu et al., 2009; He et al., 2018; Krishnamurthy et al., 2020; Laskowitz et al., 2007; Zhong et al., 2025), although clinical application of these agents in the context of AD remains to be investigated.

In addition, therapeutic strategies targeting APOE4 levels may offer promising benefits for AD (Raulin et al., 2022). Although removal of APOE4 from astrocytes or microglia has shown protective effects in amyloid and tau models (Mahan et al., 2022; Wang et al., 2021), APOE plays an essential role in cholesterol transport and lipid homeostasis systemically (Huang & Mahley, 2014). Consequently, while APOE removal reduces amyloid/tau pathology in AD mouse models (Kim et al., 2011), global ablation of APOE is associated with adverse cardiovascular outcomes and peripheral dysfunction (Ghiselli et al., 1981; Zhang et al., 1992). Rather than reducing APOE expression, modifying its conformation and enhancing lipidation state may confer benefits while minimizing systemic risk. Notably, lipid-free APOE is more prone to form aggregates with Aβ compared to HDL-associated APOE (Hubin et al., 2019). A recent study using a humanized APOE antibody targeting non-lipidated APOE showed preferential binding to pathological APOE species and reduced amyloid burden in APP/PS1/E4 mice (Liao et al., 2018). These findings further highlight the therapeutic potential of modifying APOE structure and lipidation without altering total expression, which can be accomplished by using HDL mimetic peptides including 4F.

The current study raises several important questions for future research. While we focused on astrocytes, APOE is also expressed in microglia, which play a central role in neuroinflammation and amyloid clearance. Future studies investigating the impact of 4F on APOE secretion and lipidation in APOE isoform-specific iPSC-derived microglia, as well as microglial phagocytosis and inflammatory responses, may offer additional therapeutic targets for modulating AD pathogenesis. Moreover, APOE lipidation plays a key role in promoting Aβ clearance (Verghese et al., 2013) and modulating neuroinflammation, with APOE3 isoform exhibiting enhanced lipid/cholesterol efflux and decreased neuroinflammatory responses compared to APOE4. Prior studies have shown that APOA-I overexpression or HDL mimetic peptides such as 4F can ameliorate amyloid pathology and limit neuroinflammation (Cimmino et al., 2009; Fernandez et al., 2019; Lewis et al., 2010; Lynch et al., 2003), further supporting their therapeutic potential. However, the functional effects of 4F on tau pathology, APOE4-related neurotoxicity, and neuroinflammation in iPSC-derived models or in humanized APOE mice remain to be fully elucidated.

In summary, this study provides compelling evidence that treatment with the HDL mimetic peptide 4F significantly enhances APOE secretion and lipidation, counteracts the inhibitory effect of Αβ42, and rescues APOE4-associated lipid droplet accumulation in astrocytes. These effects were observed across APOE genotypes, suggesting a shared pathway by which 4F promotes APOE function, while rescuing APOE4-related deficits. Together, these findings highlight the therapeutic potential of HDL mimetic peptides like 4F for targeting APOE4-associated lipid dysfunction and Alzheimer’s disease pathogenesis.

---

**Figure S1.**
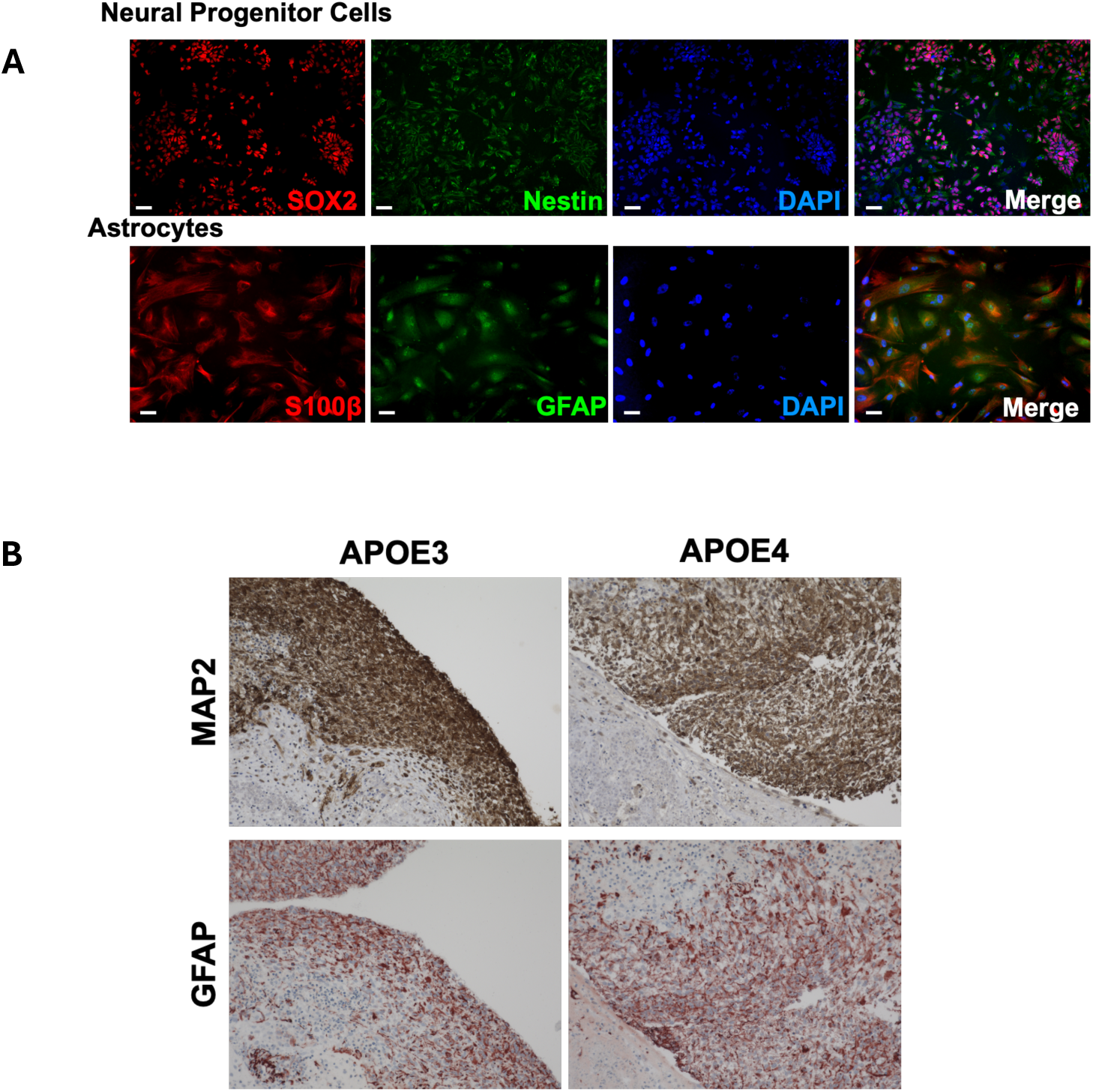
Characterization of iPSC-derived astrocyte and organoid di:erentiation. **(A)** iPSCs were di+erentiated into NPCs via dual smad inhibition, followed by di+erentiation into astrocytes for 35-40 days using astrocyte di+erentiation kit (ScienCell). Representative immunofluorescent images of NPCs expressing Sox2/Nestin and mature astrocytes expressing GFAP and S100β. **(B)** iPSCs were spontaneously allowed to di+erentiate into organoids in non adherent G-Rex (Gas-Permeable Rapid Expansion) flasks for at least 30 days. Representative immunohistochemistry images of mature cerebral organoids stained for neurons (MAP2) and astrocytes (GFAP).

## Data availability

The datasets generated and analyzed for the current study are available from the corresponding author upon reasonable request.

## Funding statement

This work was supported by the National Institutes of Health (NIH) grants (AG058081, AG081426, and AG AG077772), the University of Minnesota Office of Academic Clinical Affairs (FRD #19.29), and the College of Pharmacy of the University of Minnesota.

## Conflict of Interest Disclosure

The authors declare no conflicts of interest.

## Ethics Approval Statement

All animal used in this study were approved by the Institutional Animal Care and Use Committee (IACUC) at the University of Minnesota under protocol 2207-40221A, and performed in accordance with the university guidelines.

## Patient Consent Statement

Patient consent was not required for this study as no human subjects were used.

## Permission to Reproduce Material from Other Sources

The authors confirm that all figures, tables, and text in this manuscript are original and have not been previously published; no permission to reproduce material from other sources is required.

## Acknowledgments

We graciously thank Dr. Matthew Blurton-Jones at the University of California Irvine for providing the iPSCs used in this study and Dr. Julia TCW at Boston University for sharing the astrocyte differentiation procedures. We are also thankful to Amanda Vegoe and Dr. Timothy O’Brien from the University of Minnesota Stem Cell Institute for their assistance with generating 3D organoids. We are grateful to the staff of the Comparative Pathology Shared Resource at the University of Minnesota for their excellent technical assistance in processing the cerebral organoids used in this study.

## Notes

### Competing Interest Statement

The authors have declared no competing interest.

